# Hepatitis D virus infection of stem cell-derived hepatocytes triggers an IFN- and NFκB-based innate immune response unable to clear infection

**DOI:** 10.1101/2022.08.11.502443

**Authors:** Frauke Lange, Jonathan Garn, Holda Anagho, Thomas von Hahn, Thomas Pietschmann, Arnaud Carpentier

## Abstract

Human pluripotent stem cell-derived hepatocyte-like cells (HLCs) are a valuable model to investigate host-pathogen interactions of hepatitis viruses in a mature and authentic environment. Here, we investigated the susceptibility of HLCs to the Hepatitis D Virus (HDV), a virus that in co-infection with HBV is responsible for the most severe form of viral hepatitis. Cells undergoing hepatic differentiation became susceptible to HDV infection after acquiring expression of the Na^+^-taurocholate cotransporting polypeptide (NTCP), the receptor mediating HBV and HDV entry. Inoculation of mature HLCs with HDV lead to increasing amounts of intracellular HDV RNA and accumulation of the HDV antigen in the cells. The infection was abrogated when using known entry inhibitors targeting NTCP or by disrupting genome replication using the nucleoside analogue Ribavirin. Upon infection, the HLCs mounted an innate immune response based on induction of the interferons IFNB and L, but not IFNA, and were associated with an upregulation of interferon-stimulated genes. The intensity of this immune response positively correlated with the level of viral replication and was dependant on both the JAK/STAT and NFκB pathway activation. Importantly, neither this innate immune response nor an exogenous treatment of IFNα2b inhibited HDV replication. However, pre-treatment of the HLCs with IFNα2b reduced viral infection, suggesting that ISGs may limit early stages of infection.

This novel HDV *in vitro* infection model represents a valuable tool for studying HDV replication and investigating candidate antiviral drugs in cells displaying mature hepatic functions.

**Lay summary:** HDV can infect stem cell-derived hepatocytes through an NTCP-mediated entry process. Infection triggers an IFN and NFκB dependent innate immune response. However, viral replication seems unaffected by this innate response or by exogenous IFN treatment.

**Graphical abstract:** 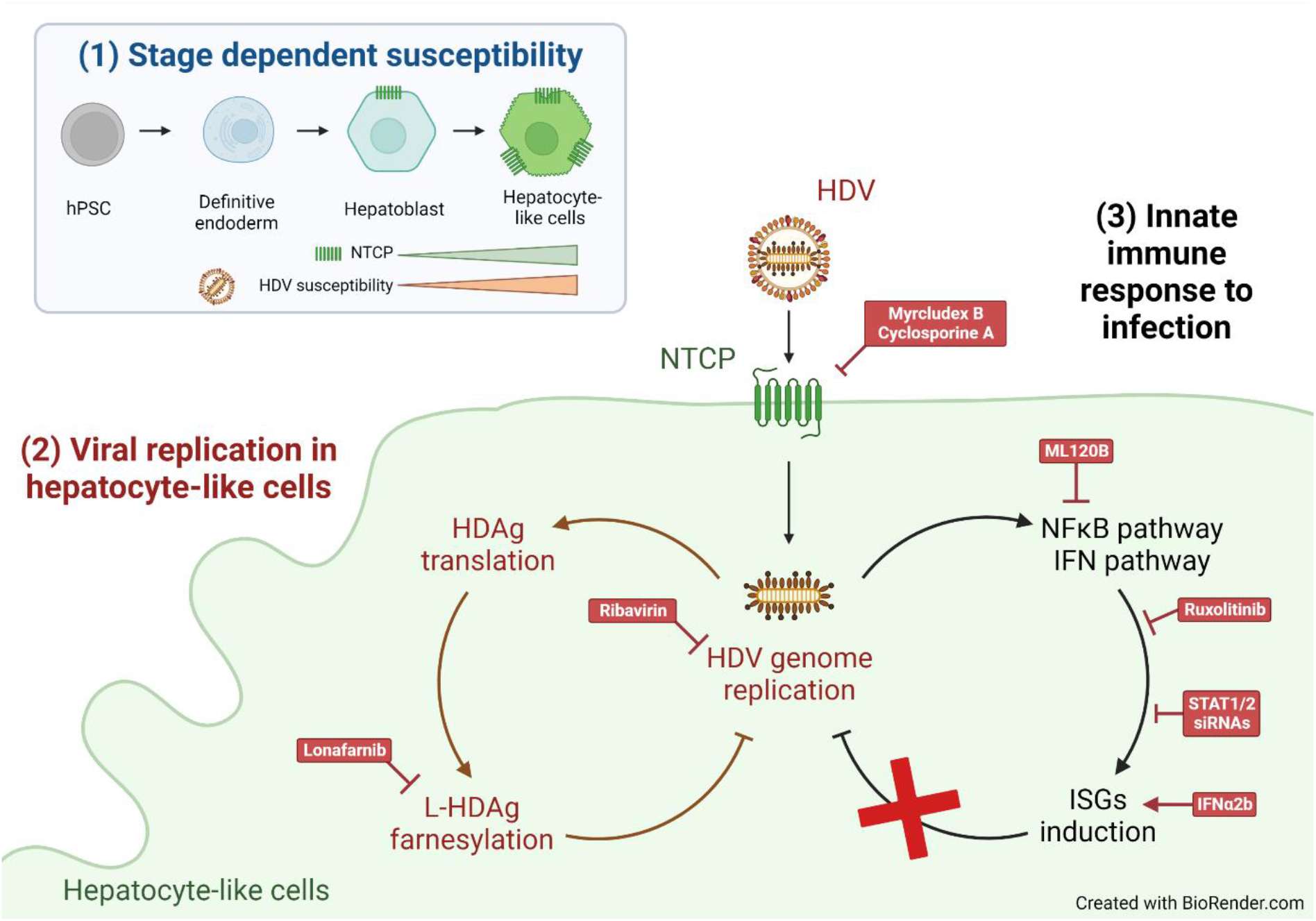

## INTRODUCTION

*In vitro* studies of hepatitis virus infection rely on cell culture models that accurately reproduce infection in the natural host: the adult hepatocyte. Though primary culture of human hepatocytes (PHHs) are considered the gold standard of *in vitro* models, their experimental application is constrained by their scarcity and donor to donor variability.^1^ Human pluripotent stem cells (hPSC)-derived hepatocyte like cells (HLCs) have shown to be a valuable alternative, reproducing key metabolic features of PHHs without the aforementioned limitations.^2^ HLCs are permissive for hepatitis virus infection both *in vitro* (HAV,^3^ HBV,^4^ HCV,^5^ HEV^6^) and *in vivo*^7^ and are being used to investigate host pathogens interactions (reviewed in ^8^), with applications ranging from reproducing polarized secretory pathways^3^ to studying single nucleotide polymorphisms-related permissivity.^9^

One aspect of viral infection that is difficult to study in a transformed cell line is the interplay between the virus and the innate immune response. Most hepatoma cell lines are notoriously immuno-incompetent when compared to PHHs. HLCs on the other hand accurately reproduce the innate immune response of PHHs. Upon HCV infection, HLCs trigger a strong interferon (IFN) based immune response that is able to clear the infection.^10^ In contrast, HBV replication in HLCs does not trigger an innate immune response, reflecting HBVs “stealthy” nature observed in PHHs.^11^

To our knowledge, no study has yet examined the susceptibility of HLCs to HDV. HDV is the smallest virus known to infect humans. It is responsible for the most severe immune driven form of viral hepatitis: Hepatitis Delta.^12^ Hepatitis Delta occurs in patients co-infected with HBV and HDV, or in chronic HBV positive patients superinfected with HDV. On its own HDV is defective: While the HDV RNA genome can replicate on its own, the virus outer layer carries only HBV surface antigens (HBsAg). Hence, HDV is dependent on HBV to enter the hepatocyte through a process involving the HBV entry receptor Na^+^-Taurocholate cotransporting peptide (NTCP, encoded by the *SLC10A1* gene),^13^ and to assemble and secrete new progeny virions. Nevertheless, HDV entry, genome replication, Hepatitis D Antigen (HDAg) translation, and post-transcriptional modification can accurately be reproduced *in vitro* in the absence of HBV.^12^

Because transformed hepatoma cell lines do not express the virus receptor NTCP, *in vitro* studies on HDV infection have long been limited to the PHHs model.^14^ To bypass this restraint, studies have artificially overexpressed NTCP as a prerequisite for investigating HBV/HDV infection. More recently, a differentiated HepaRG cell model that is produced through *in vitro* differentiation of an immortalized hepatic progenitor cell line constitutes a more relevant cell line to study HBV and HDV^15,16^. However, long and heterogeneous differentiation, poor infection efficiency, and restriction to one single host genetic background may limit its applications.

Host pathogens interactions and innate immune response to HDV infection have mostly been investigated in hepatoma cell lines (Huh7, HepG2) overexpressing NTCP and in differentiated HepaRG.^17^ PHHs have previously been demonstrated to produce a strong induction of interferons (IFNs) and interferon stimulated genes (ISGs) when infected with HDV, but only modified cell lines allowed for an in-depth study of this response. It was recently shown that in cell lines HDV RNA is sensed by the cytosolic pattern recognition receptor (PRR) melanoma differentiation-associated protein 5 (MDA5, *IFIH1* gene), but not by the retinoic acid-inducible gene I (RIG-I, *DDX58* gene) or Toll Like Receptors (TLRs). MDA5 triggering then leads to the induction of IFNB and L, but not A, and ISGs.^18, 19^ This IFN response decreases infection in dividing transformed cells,^20^ but has no antiviral effect in mature resting cells like PHHs.^17^ Moreover, exogenous *in vitro* treatment with IFNA or L showed limited effect at an early time of infection, but had no antiviral effect on established infections.^18^ This is consistent with clinical data that a IFN monotherapy only rarely leads to viral eradication in patients. Recent developments have focused on targeting the viral replication cycle instead: For example, the entry inhibitor Myrcludex B (licensed under the name “Bulevirtide”), which is a synthetic preS1-peptide derived from the L-HBsAg, prevents interaction of the virus with NTCP.^21^ Lonafarnib, a farnesyl transferase inhibitor, targets the assembly of progeny HDV.^22^

The role of the NFκB (“nuclear factor kappa-light-chain-enhancer of activated B cells”) pathway during viral infection in hepatocytes has recently been identified.^23^ In hepatic cell lines, the activation of the NFκB pathway is necessary for the full activation and maintenance of the antiviral IFN immune response after infection by various hepatotropic viruses, including HDV.^23^

Here, we describe for the first time *in vitro* infection of HLCs with HDV. HLCs support NTCP-dependent viral entry, efficient replication of the viral genome, and translation of the viral antigen. Importantly, the cells react to the infection by inducing an innate immune response dependent on both IFN and NFκB pathways. However, viral replication itself is relatively unaffected by either endogenous or exogenous IFN. This new *in vitro* model of HDV infection constitutes a promising new tool to study HDV host-pathogen interactions in the context of mature non-transformed hepatocytes.

## MATERIAL & METHODS

Detailed material and methods can be found in the Supplementary materials.

## RESULTS

### Hepatic differentiation and stage dependent susceptibility to HDV

HLCs were differentiated as previously described.^24^ Briefly, hPSCs (D0) are subjected to sequential growth factors and small drugs treatment to successively differentiate into definitive Endoderm (D4), Hepatoblasts (D8, D12), and mature HLCs (D15) (Fig 1A). RNAseq analysis of the cells at different stages of differentiation (GEO accession number: GSE132606) was used to follow the expression of known HDV host factors. Among them, the entry factors EGFR,^25^ GPC5^26^ and NTCP (*SLC10A1)* were expressed only during and after hepatic specification (Fig 1B). Inoculation of cells at different stages of differentiation confirmed susceptibility to infection only from D8 on (Fig 1C), correlating with expression of NTCP in the cells. Moreover, bypassing NTCP-mediated entry by transfecting the HDV genome into NTCP-negative hPSCs allowed viral genome replication (Sup Fig 1A), translation of the HDAg (Sup Fig 1b) and production of infectious progeny HDV if co-transfected with a plasmid encoding the HBsAg (Sup Fig 1C). Interestingly, the stage-related induction of NTCP expression is concomitant with the induction of expression of *IFIH1*, the gene coding for MDA5, the main PRR responsible for detection of HDV RNA.^18^ Replication of HDV RNA in transfected MDA5-negative hPSCs did not lead to upregulation of ISGs (Sup Fig 1D). HLCs expressed higher level of NTCP than PHHs and Huh7^NTCP^ (Fig 1D), and NTCP protein was detected in almost every HLCs (Fig 1E). Altogether, our data show that cells undergoing *in vitro* hepatic differentiation become susceptible to HDV infection when inducting the expression of the entry factor NTCP.

**Figure 1:**
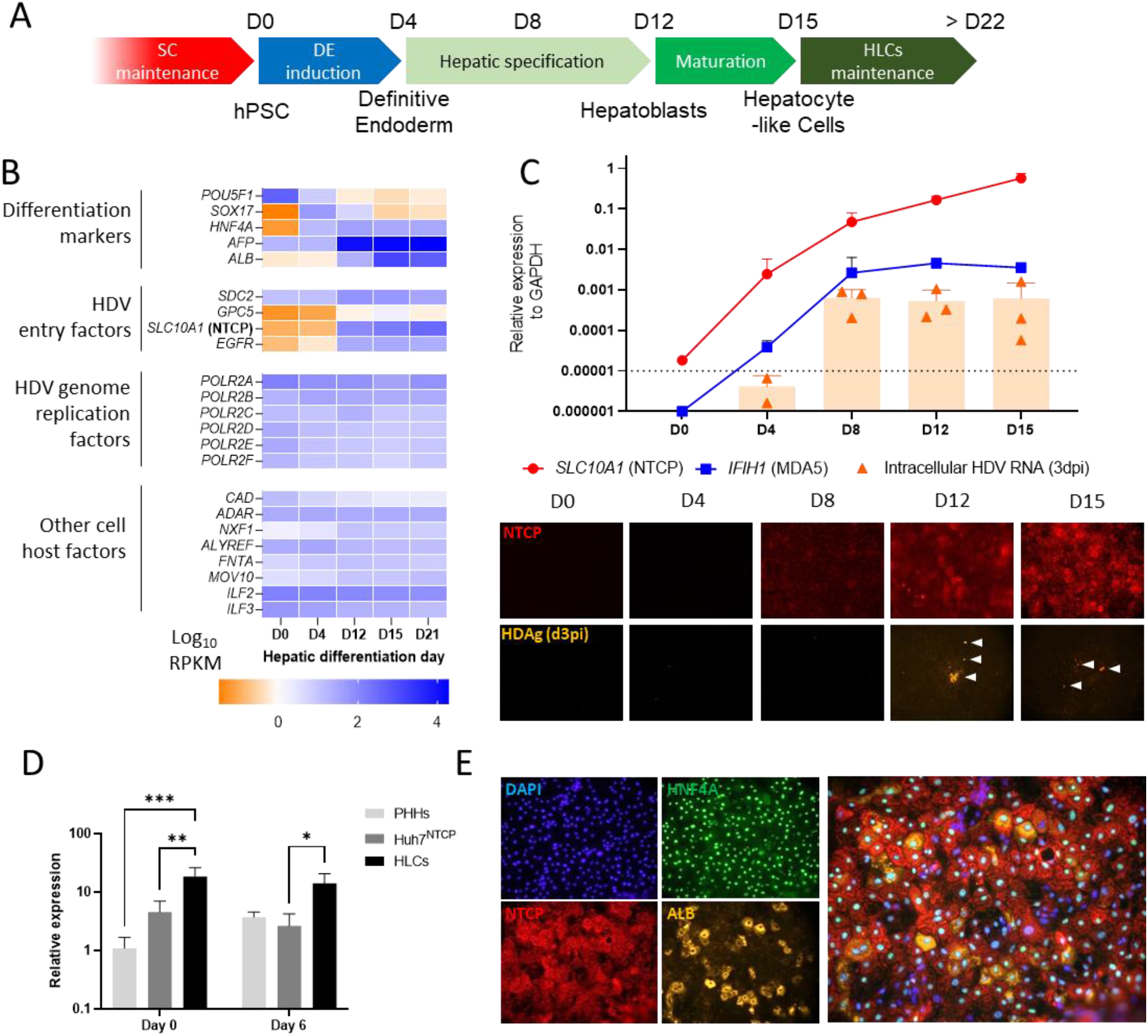
HLCs susceptibility to HDV is stage dependent and correlates with expression of NTCP. (A) Stepwise protocol of hepatic differentiation. (B) Stage-dependent expression of selected differentiation markers and known HDV host cell factors. (C) Cells at various stages of differentiation (D0, D4, D8, D12, and D15) were inoculated with HDV (n=3). 3 days later intracellular HDV RNA (assessed by RTqPCR, triangles depict individual experiments, dotted line depicts technical limit of detection) and HDAg expression (IFA, magnification 10x) were assessed and compared to expression of *SLC10A1* (NTCP) and *IFIH1* (MDA5) (lines, assessed by RTqPCR) at the time of inoculation. (D) Expression of *SLC10A1* (assessed by RTqPCR) in PHHs, Huh7^NTCP^ and HLCs at day 0 and 6 of in vitro culture, relative to PHHs at day 0. (E) Cell surface expression of NTCP and expression of hepatic markers in HLCs at the time of HDV inoculation (D15) (magnification 10x).

### Cell entry and replication inhibitors ablate HDV infection of HLCs

We inoculated HLCs with infectious HDV at a multiplicity of infection (MOI) of 0.5, for 6 hours in presence of 4% Polyethylene glycol. In the three following days, we could detect increasing amounts of intracellular HDV RNA by RTqPCR (Fig 2A), consistent with an efficient intracellular replication of the viral genome. We could also visualize by immunofluorescent assay (IFA) accumulation of HDAg in both nucleus (Fig 2B arrows) and cytoplasm (Fig 2B dotted shape) of infected HLCs. At early time points (day 1 and 3 post inoculation (dpi)), HDAg positive HLCs could rarely be visualized, while at a later time point, the intensity of staining increased, which is consistent with the accumulation of HDAg in both the nucleus and cytoplasm (Sup Fig 2A). Moreover, the absence of foci of HDAg positive cells (as seen in Huh7^NTCP^), even at later time point, is consistent with quiescence of HLCs, the lack of cell division-mediated spreading, and the inability of HDV to spread in the absence of a HBV helper virus (Sup Fig 2B). HDV infection of HLCs was efficiently blocked by Myrcludex^21^ or Cyclosporine A,^27^ which both prevent the interaction of the virus surface proteins with NTCP (Fig 2C). Intracellular replication of HDV RNA was also abrogated by treating the HLCs with Ribavirin (Fig 2C), a guanosine ribonucleoside analogue, which exhibits a strong *in vitro* antiviral action against a wide range of viruses,^28^ including HDV.^29^ In sum, our results show that HLCs support authentic viral entry, intracellular replication of the viral genome, and translation of the viral protein HDAg.

**Figure 2:**
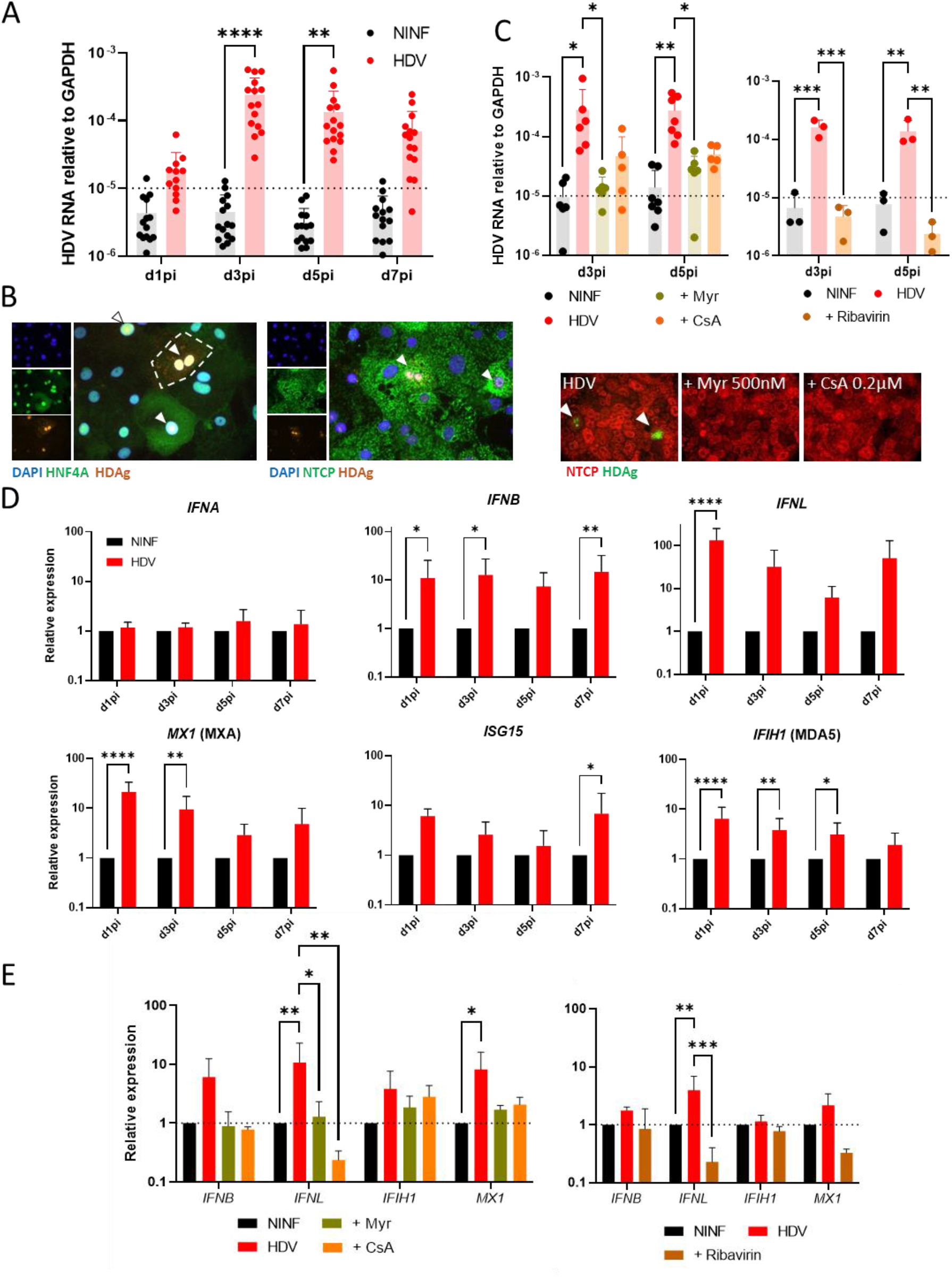
HDV infection of HLCs triggers an IFN based innate immune response. (A) Intracellular HDV RNA assessed by RTqPCR, from day 1 and every 2 days post inoculation with HDV (MOI 0.5) (n=15). Intracellular HDV RNA titers are depicted with dots and bar representing respectively individual experiments and mean; and are compared to technical detection level (dotted line) (B) Intracellular HDAg expression (arrowhead) detected by IFA (magnification 100X) in HLCs, at day 5pi. (C) Myrcludex (Myr, 500nM, n=6), Cyclosporine A (CsA, 0.2μM, n=6) and Ribavirin (0.5mM, n=3) pre-treatment and inhibition of HDV infection, assessed by quantification of intracellular HDV RNA and staining for HDAg (Magnification 20x). (D) Induction of expression of IFNs and ISGs upon HDV infection of HLCs, relative to non-infected cells (NINF) at the same time point. Cellular genes RTqPCR results are illustrated as average value of n=6-12 independent experiments (E) Myrcludex (Myr, 500nM), Cyclosporine A (CsA, 0.2μM) and Ribavirin (0.5mM) pre-treatment and inhibition of HDV dependent induction of immune response (d5pi).

### Strategies to improve HLCs susceptibility to HDV

Overall, inoculation of HLCs at MOI 0.5 led to a low percentage of detectable infected cells (estimated around 2%). HLCs are less permissive to infection than Huh7^NTCP^, as visualized by HDAg staining (Sup Fig 2B) and RTqPCR for intracellular HDV RNA (Sup Fig 2C). The majority of HLCs express NTCP (Fig 1E), but the receptor may not be accessible to the virus due to cell polarization. Disruption of the tight junctions by EDTA treatment has been shown to improve viral access to NTCP in polarised cell lines and increase infection with HBV.^30^ In our HLCs, EDTA and EGTA pre-treatment disrupted tight junctions and improved the NTCP-mediated uptake of indocyanin green (Sup Fig 3A).^31^ However, the pre-treatment did not significantly enhance intracellular HDV RNA amounts (Sup Fig 3B) and HDAg positive cells were only increased by a slight percentage (Sup Fig 3C).

The limited number of HDAg positive cells may also be due to inhibition of viral replication within the cells. Farnesylation of the large isoform of the HDAg has been described to inhibit viral genome replication.^32,33^ Inhibiting HDAg farnesylation using Lonafarnib improves HDV genome replication in hepatoma cell lines.^34^ In our HLCs, Lonafarnib treatment increased both intracellular HDV RNA amounts (Sup Fig 4A) and percentage of HDAg positive cells detected by IFA significantly (Sup Fig 4B). Remarkably, a Lonafarnib treatment led to a higher HDAg immunofluorescent signal concentrated only within the nucleus of infected cells (Sup Fig 4B, arrows).

These data show that, although the presence of a restriction mechanism at entry level cannot be excluded, the intracellular accumulation of farnesylated L-HDAg significantly inhibits viral replication of HDV RNA in the infected HLCs.

### Cellular innate immune response to HDV infection

HLCs accurately react to cytoplasmic PRR triggering, using Poly(I:C) or upon HCV infection, producing IFNs and upregulating antiviral ISGs.^10^ In order to assess this IFN-based innate immune response upon HDV infection, inoculated HLCs were harvested at various dpi and compared to naïve cells at similar time points. Upon infection, type 1 interferon *IFNB* mRNA was significantly upregulated as opposed to *IFNA* mRNA (Fig 2D). Notably, type III *IFNL* mRNA was strongly over expressed in the first dpi, and then decreased over days. Canonical ISGs *MX1* (MXA), *ISG15* and *IFIH1* (MDA5) were induced throughout the experimental time line (Fig 2D). Levels of induction of IFNs and ISGs correlated with the level of intracellular HDV RNA, particularly at d3 and d5pi (Sup Fig 5). Myrcludex, CsA or Ribavirin treatment prevented the induction of these immune genes (Fig 2E), confirming that the triggering of the HLCs’ immune response is dependent upon authentic HDV infection and viral genome replication. Interestingly, Lonafarnib treatment of HDV infected HLCs, which allowed higher levels of viral genome replication (Sup Fig 4A), led to increased levels of innate immune activation (Sup Fig 4C), further confirming that the innate immune response observed is triggered by the viral replication.

Our results show that HDV infection triggers an innate immune response dependent on and correlating with the level of viral replication.

### HDV innate immune response is dependent upon activated JAK/STAT and NFκB pathways

Triggering of HLCs’ cytoplasmic PRRs (RIG-I and MDA5) using Poly(I:C) transfection leads to nuclear translocation of Interferon regulatory factor 3 (IRF3) and induction of expression of IFNs and canonical ISGs, consistent with an activation of the JAK/STAT pathway (Sup Fig 6A). PRR triggering also leads to nuclear translocation of the NFκB subunit RelA (also known as p65) and upregulation of the NFκB target gene Tumour Necrosis Factor Alpha-Induced Protein 3 (*TNFAIP3*, also known as A20),^35^ consistent with activation of the NFκB pathway (Sup Fig 6B). HDV infection of HLCs also induced expression of *TNFAIP3*, however only in a limited and transient way, but consistent with an activation of the NFkB pathway (Sup Fig 6C). In order to assess the role of both IFN and NFκB pathways upon HDV infection, we treated the HLCs with specific inhibitors. Ruxolitinib (Fig 3, green), a specific JAK1 and 2 inhibitor blocking the IFN pathway,^10^ and ML120B (Fig 3, blue), a specific IKKβ inhibitor blocking the NFκB pathway,^23^ both abrogated the induction of IFN and canonical ISG in HDV infected HLCs (Fig 3A). However, none of these treatments showed a positive effect on intracellular HDV RNA (Fig 3B) and HDAg expression (Fig 3C). Induction of innate immune response could also be inhibited by transfecting the cells with siRNAs targeting STAT1 and STAT2 (Fig 4A, B), but had no significant effect on viral replication (Fig 4C). HDV infection of the hepatoma cell line Huh7^NTCP^ did not induce a significant innate immune response; and treatment with inhibitors or Lonafarnib did not change the level of viral genome replication in this model (Sup Fig 7A, B). It has been hypothesized that the HDV ribozyme may activate PKR.^36^ In order to assess the role of PKR in infected HLCs, we treated the cells with the oxindole/imidazole derivative C16 PKR inhibitor.^37^ C16 treatment had no effect on the induction of the innate immune response (Sup Fig 8A) and did not modify the level of viral replication (Sup Fig 8B).

**Figure 3:**
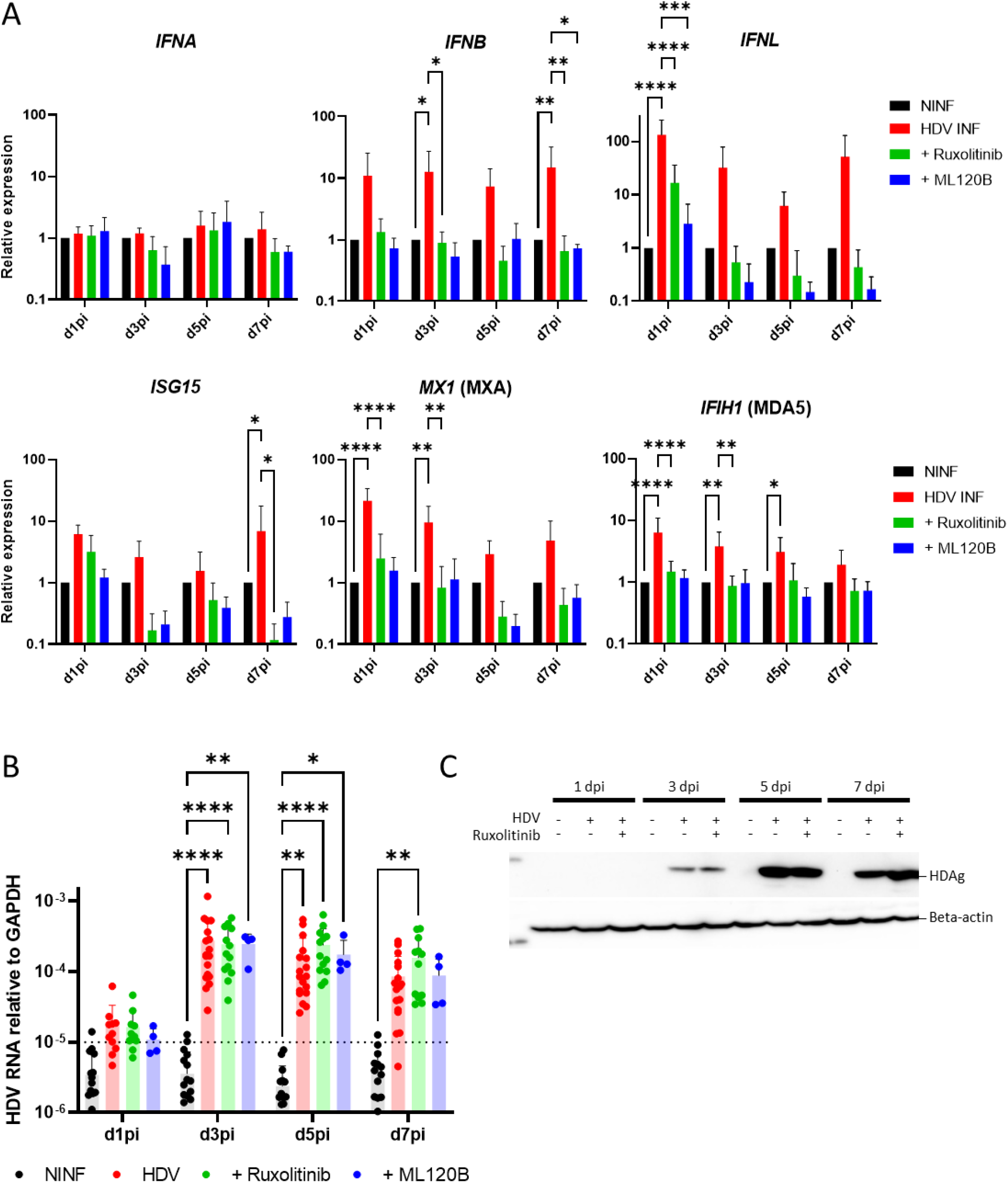
Pharmaceutical inhibition of the innate immune response does not significantly improve level of HDV replication in HLCs. Control HLCs (n=16) or HLCs treated with 10μM JAK/STAT pathway inhibitor Ruxolitinib (n=12) or 10μM NFκB inhibitor ML120B (n=4) were inoculated with HDV (M.O.I. 0.1). (A) Intracellular HDV RNA and (B) protein expression of HDAg assessed by western blot, in ctrl HLCs, HDV inoculated HLCs and Ruxolitinib treated HDV infected HLCs. (C) induction of IFNs and ISGs, assessed by RTqPCR from day 1 and every 2 days post inoculation and compared to ctrl HLCs.

**Figure 4:**
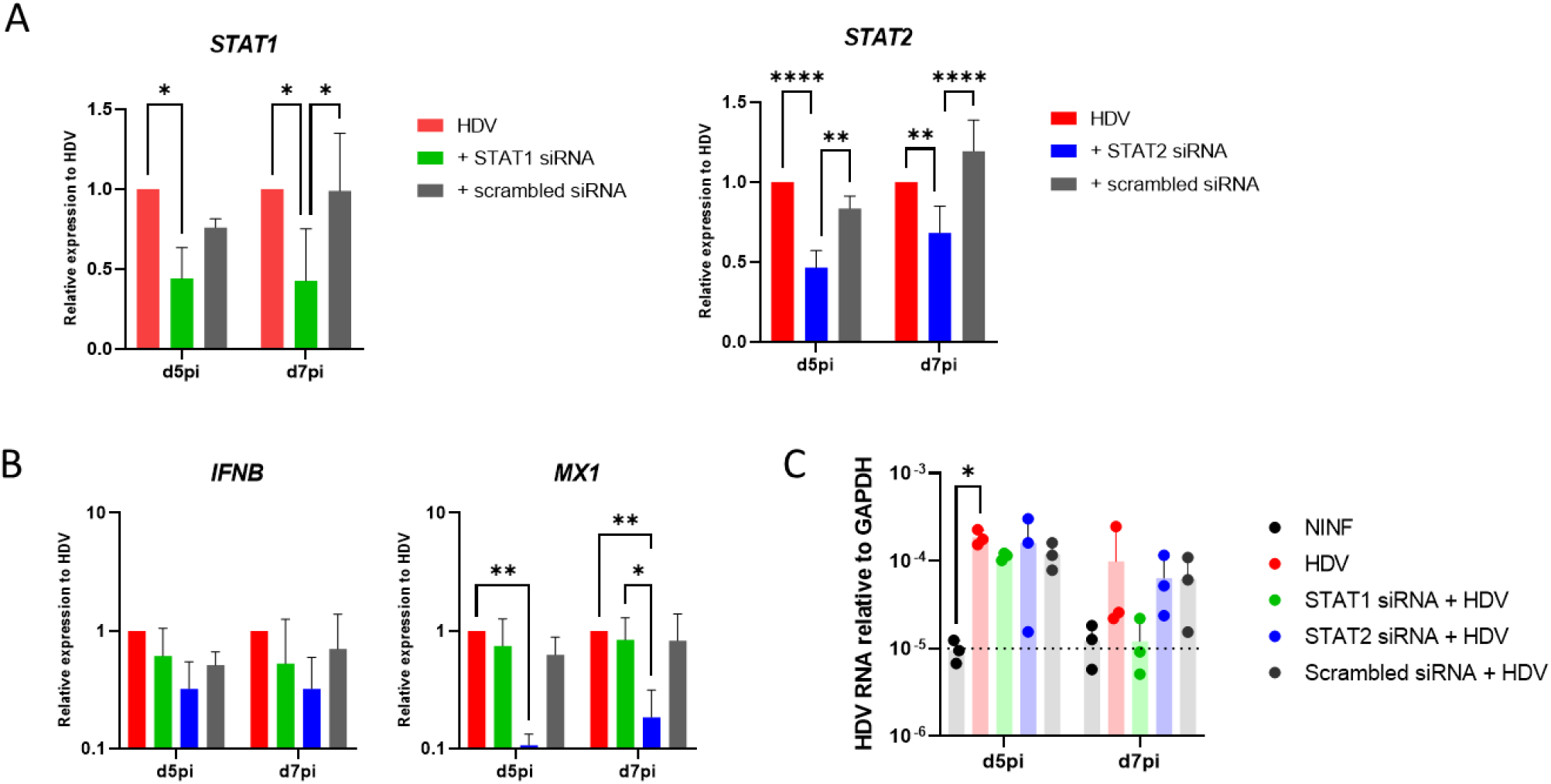
STATs are necessary for the HDV-mediated activation of the IFN and JAK STAT pathway. (A) HLCs were transfected with siRNA targeting STAT1 or 2, and effect of these siRNA was confirmed by RTqPCR quantification of the target genes expression (n=3). (B) Induction of IFN and ISGs was measured by RTqPCR in these cells. (C) Ctrl and siRNA treated HLCs were then inoculated with HDV and intracellular HDV RNA was then assessed by RT qPCR at d5 and d7pi (n=3).

Altogether, our results show that upon HDV infection, immune competent HLCs mount an IFN and NFκB-dependent, PKR-independent innate immune response. However, this antiviral immune response is not able to restrict viral replication.

### Effect of exogenous IFNα2b on HLCs permissivity to HDV

To explore the antiviral activity of exogenous IFNs, we treated HLCs, which were inoculated 3 days earlier with HDV, with 1000 IU/ml of IFNα2B (Fig 5A, B). 2 days post IFN treatment, we confirmed that IFNα2B could induce the expression of ISGs in naïve HLCs (Fig 5B, blue) and could slightly increase the induction of ISGs in HDV infected HLCs (Fig 5B, purple vs red). However, IFNα2b treatment showed no antiviral effect (Fig 5A, red vs purple).

**Figure 5:**
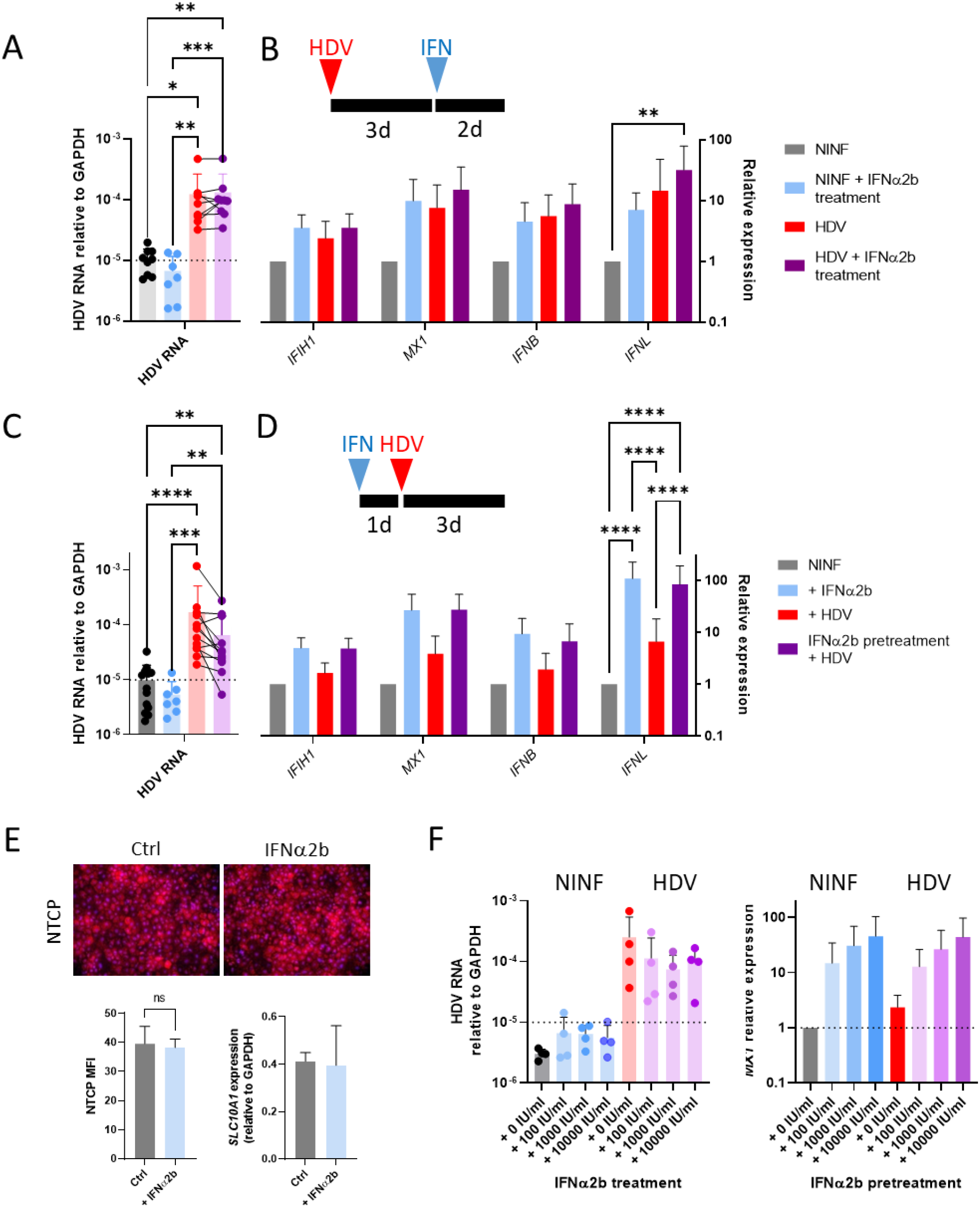
IFNα2b treatment does not reduced established viral replication, but pre-treatment partially restricts HDV infection of HLCs. (A-B) Ctrl HLCs and HLCs inoculated with HDV 3 days earlier were treated or not with 1000 IU/mL of IFNα2b (n=8). 2 days later, (A) intracellular HDV RNA, and (B) IFNs and ISGs expression were assessed by RTqPCR, compared to control untreated non-infected HLCs. (C-D) HLCs, pre-treated or not for 16h with 1000 IU/mL of IFNα2b, were inoculated with HDV. Intracellular HDV RNA (C), IFNs and ISGs expression (D) were then assessed by RTqPCR 3 days post inoculation. (E) Effect of IFNα2b on protein expression of NTCP (magnification 10x) and gene expression of SLC10A1 (NTCP). (F) Dose effect of IFNα2b as pre-treatment on HDV replication and canonical ISG (*MX1*) induction (n=4).

Pre-treating HLCs with 1000 IU/ml of IFNα2b 16 hours before inoculation with HDV allowed a mild viral restriction effect (Fig 5C, red vs purple). A pre-treatment with IFN showed a higher level of induction of ISGs and IFNs than an HDV infection (Fig 5D, blue vs red). Moreover, the IFNα2b-induction and the HDV-mediated immune activation were not additive (Fig 5C, blue vs purple). This observed viral restriction effect was not associated with a down regulation of NTCP, as assessed by IFA and RTqPCR (Fig 5E). Interestingly, IFNα2b exhibited a dose-dependent effect on the induction of canonical ISG MX1 but the observed antiviral effect could not be increased when using a higher dosage (Fig 5F). Collectively, our data show that IFNα2b is not able to inhibit viral replication in immunocompetent infected HLCs. However, IFNα2b treatment renders naïve cells more resistant to viral infection.

## DISCUSSION

Here we describe for the first time that stem cell-derived hepatocytes, also known as HLCs, are permissive to HDV infection *in vitro*. HLCs support authentic NTCP-mediated viral entry, viral genome replication and translation of HDAg (Figure 2). Moreover, we show that HLCs become susceptible to HDV by acquiring NTCP expression during hepatic specification. Intriguingly, as the cells start expressing NTCP and become permissive to HDV, they start expressing the PRR MDA5, suggesting they can then detect replicating viral RNA using PRRs instead of relying on the hPSCs’ constitutive immunity.^38^

Our model is based on the HDV mono-infection of HLCs, making this model safer to use and more accessible than models based on co-infection with HBV, a virus notoriously difficult to produce *in vitro*.^39^ Though in the absence of HBV, our model does not allow for assembly and production of progeny viruses, and infection cannot spread to adjacent cells, our HLCs monoinfection model brings valuable insights into the host-pathogen interactions of HDV. This is of particular interest, when studying the innate immune response of an HDV infection. Because HBV infection does not raise IFN and ISGs in hepatocytes,^40^ the innate immune response observed in co-infected patients is hypothesized to be driven by HDV (reviewed in ^17^). Accordingly, we previously showed that HBV infection does not trigger any innate immune response in HLCs.^4,11^ In our present study we show that, on the other hand, upon HDV monoinfection, HLCs mount a strong innate immune response. The induction of IFNL and IFNB, but not IFNA (Fig 2D), is consistent with what has been described in differentiated HepaRG and in PHHs.^18,19^ Importantly, this response was dependent on proper viral entry and genome replication (Fig 2E), and correlated with the level of intracellular viral replication (Sup Fig 5). Significant induction of these immune genes could be detected despite a low percentage of infected cells (Sup Fig 2A). When using a higher MOI (up to 5 infectious units per cell), we could obtain a marginal increase of the intracellular HDV RNA titer and of the percentage of HDAg positive cells (Sup Fig 9A, B), but no significant effect was seen on induction of IFNs and ISGs (Sup Fig 9C). Here, we could decipher some aspects of the innate immune response after HDV inoculation. We describe that upon HDV infection, the induction of innate immune genes in HLCs is not only under the control of the JAK/STAT pathway, similar to after infection by other hepatotropic RNA viruses like HCV;^10^ but also relies upon the activation of the NFκB signalling cascade (Fig 3). In HepaRG cells, HDV has been shown to induce NFκB driven genes.^23^ Similarly, an HDV infection of HLCs transiently elevates the expression of NFκB driven genes (Sup Fig 6C), and abrogation of the NFκB signalling cascade critically hampers the induction of the IFN-based innate immune response (Fig 3).

Moreover, we show that viral replication is not affected by the triggered innate immune response: Abrogating either JAK/STAT or NFκB pathway efficiently blocks ISGs induction but did not increase viral replication (Fig 3). Recently, inducers of the NFκB pathway have been shown to reduce HDV replication in immune competent PHHs and HepaRG.^41^ The HLCs monoinfection model offers the ideal platform to confirm if increased NFκB activation would lead to viral inhibition and whether it is dependent on ISGs induction.

Importantly, we show that exogenous IFNα2b treatment had no effect on established HDV infection of immunocompetent HLCs (Fig 5A). These findings are consistent with clinical observations that IFN treatment alone rarely leads to viral clearance (reviewed in ^17^). Interestingly, when pre-treating with IFNα2b, the HDV infection of HLCs was partially restricted (Fig 5C). This suggests that IFNα2b treatment may render naïve hepatocytes less susceptible to infection by inducing expression of ISGs. In settings where HDV infection can spread (i.e. in presence of HBV), IFN treatment could limit the level of viral replication by decreasing the number of secondary infection events. Recently, Zhang et al. showed that exogenous IFNs exert a strong antiviral effect in non-quiescent cells by destabilizing the viral RNA during the cell division process when viral structures, usually sequestered in the nucleus, are transiently released into the cytoplasm during metaphase and anaphase.^20^ It is unclear whether the antiviral mechanisms observed in our IFN-treated quiescent HLCs are the same, targeting HDV RNA on their way from the cell surface to the nucleus. Alternatively, viral restriction could occur earlier at the entry/membrane fusion steps, and for example involve IFITMs, a family of ISGs restricting viral entry, including in hepatocytes.^42^

A low percentage of infected cells suggests the existence of restriction mechanisms against HDV infection in HLCs. It has been shown that tight junctions of hepatoma cell lines restrict the accessibility of HBV to NTCP.^30^ In our HLCs, disrupting the tight junctions did not have a strong positive effect on intracellular HDV RNA or percentage of HDAg positive cells, suggesting that access to NTCP is not the most restricting factor. Recently, a study on hepatoma cell lines showed that treating cells with the farnesyl transferase inhibitor Lonafarnib improved viral genome replication by inhibiting the formation of farnesylated HDAg exhibiting inhibitory effect on HDV genome replication.^34^ In HLCs, Lonafarnib treatment has a similar effect, leading to higher viral genome replication, increased number of positive cells, and concentration of HDAg in the nucleus, where viral genome replication takes place. These findings suggest that a feedback loop effect on the viral genome replication by accumulated farnesylated HDAg may partially explain the low percentage of detectable infected cells. However, other restriction mechanisms cannot be excluded. Recently, new IFN-independent mechanisms of viral restriction have been described in hepatocytes: Among them, the IRF1 constitutive immunity^43^ and the lectin CD302 driven restriction^44^ have been associated with a resistance to infection by other RNA hepatitis viruses.

While under expressed in hepatoma cell lines, HLCs are able to express these restriction factors at a level similar to PHHs, making HLCs a relevant model to investigate anti-HDV restriction.

## List of abbreviations

CsA: Cyclosporine A
EDTA: Ethylenediaminetetraacetic acid
EGFR: Epidermal growth factor receptor
EGTA: ethylene glycol-bis(β-aminoethyl ether)-N,N,N′,N′-tetraacetic acid
GPC5: Glypican 5
HAV: Hepatitis A Virus
HBV: Hepatitis B Virus
HBsAg: HBV surface antigen
HCV: Hepatitis C Virus
HDAg: HDV antigen
HDV: Hepatitis D Virus
HEV: Hepatitis E Virus
HLC: hepatocyte-like cell
hPSC: Human Pluripotent Stem Cell
IFIH1: interferon induced with helicase C domain 1
IFITM: Interferon-induced transmembrane protein
IFN: Interferon
IKK: inhibitor of nuclear factor kappa-B kinase
iPSC: induced Pluripotent Stem Cells
IRF: Interferon regulatory factor
ISG: interferon-stimulated gene
IU: International unit
JAK: Janus kinase
MDA5: melanoma differentiation-associated protein 5
MOI: multiplicity of infection
MX1: MX Dynamin Like GTPase 1
NFkB: Nuclear factor kappa-light-chain-enhancer of activated B cells
NTCP: Na^+^ taurocholate cotransporting polypeptide
PHH: Primary human hepatocyte
PRR: pattern recognition receptor
RIG-I: retinoic acid-inducible gene I
STAT: Signal transducer and activator of transcription protein
TLR: Toll like receptor
TNFAIP3: Tumour Necrosis Factor Alpha-Induced Protein 3

## Acknowledgements

We would like to thank Prof. Heiner Wedemeyer (MHH) for sharing Myrcludex with us and are grateful to Sonja T. Jesse for editing the final manuscript. Thank you to all the members of the Institute for Experimental Virology for their support.

## Ethical and/or legal aspects

This project has been approved by the Zentrale Ethik-Kommission für Stammzellenforschung of the Robert Koch Institute (RKI), and fulfil all legal requirements according to the German Stem Cell Act / Stammzellgesetz (StZG): Genehmigung Nummer 173, 07.10.2021.

