## Supplementary data for "Hepatitis D virus infection of stem cell-derived hepatocytes triggers an IFN- and NFκB-based innate immune response unable to clear infection"

[1]: Institute for Experimental Virology, TWINCORE Centre for Experimental and Clinical Infection Research, a joint venture between Medical School Hannover (MHH) and Helmholtz Centre for Infection Research (HZI), Hannover, Germany.

[2]: Institut für Molekularbiologie, Medizinische Hochschule Hannover, Hannover, Germany, Klinik für Gastroenterologie, Hepatologie und Endokrinologie, Medizinische Hochschule Hannover, Hannover, Germany.

[3]: German Centre for Infection research (DZIF), Partner site Hannover-Braunschweig, Hannover, Germany.

[4]: Cluster of Excellence RESIST (EXC 2155), Hannover Medical School, Hannover, Germany.

http://orcid.org/0000-0002-6994-4689

### SUPPLEMENTARY MATERIAL & METHODS

#### Hepatic differentiation of hPSC

Hepatic differentiation of hPSC was described before.[24] hPSCs adapted to monolayer[S1] are cultured in mTeSR+ (Stem Cell Technologies) on plate coated with 0.4 mg/mL of growth factor reduced (GF-) Matrigel (Corning), at 37°c and 21% oxygen. For passage, cells are treated with Accutase for a few minutes at 37°c, and then cultured in mTeSR+ medium in presence of 10µM Y-27632 Rho Kinase Inhibitor. Definitive endoderm (DE) differentiation was induced using the STEMdiff™ Definitive Endoderm Kit, according to the manufacturer's instruction. 2.10^6 hPSCs seeded on low concentrated Matrigel (dilution 1 in 80 in DMEM/F12) were subjected to sequential treatment and DE induction was confirmed 4 days later by staining for SOX17 and FOXA2, and disappearance of the pluripotency marker OCT4. To initiate hepatic specification, DE cells were passed 1 in 3 on low concentrated Matrigel and cultured for 8 days in 45% High glucose DMEM - 45% F12 Supplement - 10% Knock-Out Serum replacement (KOSR), in presence of 1% Penicillin and Streptomycin, 1% non-essential amino acids and 1% Glutamine (Called Differentiation medium). The medium was replaced daily and supplemented with 100ng/ml of HGF and 1% DMSO. Hepatic maturation was then achieved by culturing the cells in Differentiation medium with 10^-7 M Dexamethasone. Differentiated HLCs were then maintained up to 11 days in complete WEM medium (William's E medium, 8.4% FCS, 1% Penicillin Streptomycin, 1µg/mL human Insulin, 5µg/mL Hydrocortisone 21-hemisuccinate, 1.8% DMSO) as described previously.

#### HDV production, concentration and titration

Infectious HDV were produced as previously described.[16, S2] Briefly, NTCP-overexpressing Huh7[NTCP S3] were transduced with equal amount of the plasmids pT7HB2.7, encoding the HBV surface proteins (genotype D), and pSVLD3, containing three copies of the HDV genome (genotype 1), using FuGENE® transfection reagent (3.5:1 Fugene to DNA ratio). Cells were then maintained in DMEM medium containing Blasticidine and 10% (during first 6 days) or 3% FCS. At days 8, 10 and 13 post transfection, supernatant were collected and stored at 4°c. At the end of the production phase, supernatants containing HDV were pooled together and concentrated 10 times using Amicon® Stirred Cells. Concentrated inputs were then titrated on Huh7$^{NTCP}$. 2500 cells per well of a 96w plate were incubated for 6 hours with serial dilution of inputs, in presence of 4% PEG. After 2 PBS washes, the cells were kept in culture for 5 days in DMEM + FCS + 10µg/ml Blasticidin. 5 days later, cells were stained for the Hepatitis D Antigen (HDAg) and number of positive cells foci were counted on an Olympus IX81 inverted fluorescent microscope.

**HDV inoculation and treatment**

Target cells, HLCs or Huh7$^{NTCP}$ were inoculated with HDV, respectively in complete WEM medium or complete DMEM medium 10% FCS, in presence of 4% Poly Ethylene Glycol (PEG). 6 hours later, the medium was removed and the cells extensively washed with PBS. If needed, the cells were pre-treated with various drugs for 16 hours before inoculation:

Myrcludex – Bulevirtide, a kind gift from Prof. H Wedemeyer, MHH – 500nM final

Cyclosporin A – Sigma-Aldrich, 30024-25mg – 0.2 mM final

JAK/STAT inhibitor Ruxolitinib - AdipoGen Life Sciences, AG-CR1-3624 – 10µM final

NFκB inhibitor ML120B – Merck #SML1174 - 10µM final

PKR inhibitor, oxindole-imidazole C16 – Sigma, I9785 - 200nM final

Lonafarnib – BLD Pharm, BD136650 - 0.1µM final

IFNα2b – Sigma Aldrich, SRP4595; or INTRON®A, Merck - 100 to 10000 IU/ml final

Ribavirin – Sigma Aldrich R9644-10MG – 0.5mM final

Myrcludex and Cyclosporin A were added to the medium only before and during inoculation with HDV. Ruxolitinib, ML120B, C16, Lonafarnib, IFNα2B and ribavirin were used before, during and after inoculation.

**ICG EDTA EGTA treatment**

HLCs were cultured at 37°c in presence of 1mg/ml of Indocyanin Green. 1 hour and half later, medium containing ICG was removed and the cells were washed 5 times with PBS. Pictures of the cells were acquired using an EvosFL microscope. To disrupt tight junctions, HLCs were incubated for 30 minutes in presence of 800µM EDTA or EGTA, followed by ICG incubation or HDV infection in presence of EDTA/EGTA.

**Poly(I:C) transfection**

HLCs were incubated for 24 hours with 1µg/mL of Poly(I:C) (from InvivoGen, reference: tlrl-pic) premixed with Lipofectamine 2000 (2µl Lipofectamine 2000 per 1µg Poly(I:C), 15 minutes at RT in OptiMEM). After 3 and 5 days, total RNA were isolated or cells were fixed using PFA 3% for immunostaining.

**RTqPCR**

Total cellular RNA was isolated using the NucleoSpin RNA kit (Macherey-Nagel). 250 ng of total RNA was then reverse transcribed using the Takara Reverse Transcription (RT) kit. 100 ng of cDNA per reaction were used for quantification using Takara's SYBR Premix Ex Taq II according to the manufacturers' instructions. Samples were then run on a Light Cycler 480 (Roche). Relative expression was calculated following the 2-DDCT methods.[S4] Relative intracellular HDV RNA titer was assessed using an HDV RT-qPCR protocol described earlier.[19]

**Table primers**

| | | |
|---|---|---|
| *GAPDH* | S | GAAGGTGAAGGTCGGAGTC |
| | AS | GAAGATGGTGATGGGATTTC |
| HDV RNA | S | CGGGCCGGCTGTTCTTCT |
| | AS | AAGGAAGGCCCTCGAGAACA |
| *IFIH1* (MDA5) | S | TCGAATGGGTATTCCACAGACG |
| | AS | GTGGCGACTGTCCTCTGAA |
| *IFNA* | S | GCTTTACTGATGGTCCTGGTGGTG |
| | AS | GAGATTCTGCTCATTTGTGCCAG |
| *IFNB* | S | ATGACCAACAAGTGTCTCCTCC |
| | AS | GGAATCCAAGCAAGTTGTAGCTC |
| *IFNL* (IL28) | S | CTTTAAGAGGGCCAAAGATGC |

| | | |
|---|---|---|
| | AS | CCAGCTCAGCCTCCAAAG |
| *ISG15* | S | CGCAGATCACCCAGAAGATCG |
| | AS | TTCGTCGCATTTGTCCACCA |
| *MX1* (MXA) | S | GTTTCCGAAGTGGACATCGCA |
| | AS | CTGCACAGGTTGTTCTCAGC |
| *SLC10A1* (NTCP) | S | AAGGACAAGGTGCCCTATAAAGG |
| | AS | TTGAGGACGATCCCTATGGTG |
| *STAT1* | S | CAGCTTGACTCAAAATTCCTGGA |
| | AS | TGAAGATTACGCTTGCTTTTCCT |
| *STAT2* | S | GAGCCAGCAACATGAGATTGA |
| | AS | GCCTGGATCTTATATCGGAAGCA |
| *TNFAIP3* (A20) | S | TCCTCAGGCTTTGTATTTGAGC |
| | AS | TGTGTATCGGTGCATGGTTTTA |

**Immunofluorescent staining**

For immunofluorescence assays (IFA), HLCs were fixed using PBS 3% paraformaldehyde for 30 minutes, permeabilized using PBS 0.5% Triton for 15 minutes, and blocked using PBS 3% BSA for 15 minutes. Primary antibodies (see list below) were diluted in PBS 0.1% Triton 1% BSA and incubated on cells for 1 night. After 3 washes with PBS 0.1% Triton, secondary antibodies, diluted 1 in 10000 in PBS 0.1% triton 1% BSA, were incubated for 45 minutes on cells.

| | | |
|---|---|---|
| Alpha fetoprotein (AFP) | Sigma Aldrich, A8452 | 1:330 dilution |
| Albumin | Cedarlane, CL2513A | 1:330 dilution |
| Hepatocyte Nuclear factor 4A (HNF4A) | Santa Cruz, sc-6556 | 1:100 dilution |
| Interferon regulatory factor 3 (IRF3) | Cell Signaling, #11904 | 1:400 dilution |
| p65 (RELA) | Cell Signaling, #8242T | 1:500 dilution |
| NTCP | Thermo Fisher, PA5-80001 | 1:500 dilution |
| OCT4 | Santa Cruz, sc-9081 | 1:400 dilution |

The anti-genotype 1 HDAg monoclonal antibody, named HDAg#280, was generated by Synaptic Systems GmbH (Göttingen, Germany).[S2] The HDAg#280 antibody was used at 1:500 dilution.

Images were acquired on an EVOS® FL Cell Imaging System or an Olympus IX81 inverted microscope.

**Western Blot**

Total cell population was lysed in cold RIPA buffer containing protease (Thermo Scientific) and phosphatase (Sigma Aldrich) inhibitors, and 1U/µl Benzonase Nuclease (Merck Millipore). 7.5µg of protein in PBS SDS buffer were denatured at 98°c for 5 minutes and ran on a gel electrophoresis. Proteins were transferred to a PVFD membrane, blocked and incubated with primary antibodies (HDAg#280, 1:2000; anti-β-Actin, Sigma, A3854, 1:5000) overnight at 4°c. A secondary antibody (anti-mouse HRP-coupled, 1:10.000) was applied for 1 h at room temperature. Signal was detected with SuperSignal™ West Femto Maximum Sensitivity Substrate (Thermo Scientific) or Amersham™ ECL™ Prime Western Blotting Detection Reagent (GE Healthcare) on an Intas ChemoStar Professional Imager.

**STATs siRNA assay**

siRNA against human STAT1 and STAT2 (Ambion) were used to transfect HLCs 1 day before infection, using 3µl RNAi MAX Lipofectamine and 20pmol of siRNA (40µM final) per well of a 24 well plate, following the manufacturer's instructions. Transfection was then repeated on infected cells at day 2 and 4pi.

| Target | Ambion ID | Sequence |
|---|---|---|
| Human STAT1 | s279 | UCCGCAACUAUAGUGAACCdAdG |
| Human STAT2 | s13528 | UUAGAGACCACAAUGAGCCdTdG |

**Statistics**

Statistics analyses were performed using GraphPad Prism software. 2 way ANOVA or Kruskal-Wallis tests were performed depending on data set. Correlation coefficients were assessed using a Pearson r test. Results are displayed as follow: * P value <0.05; ** P value <0.01; *** P value < 0.001. **** P value <0.0001.

**SUPPLEMENTARY REFERENCES.**

[S1] Chen KG, et al. Non-colony type monolayer culture of human embryonic stem cells. Stem Cell Res. 2012 Nov; 9(3): 237–248.

[S2] Buchmann B, Dohner K, Schirdewahn T, Sodeik B, Manns MP, Wedemeyer H, et al. A screening assay for the identification of host cell requirements and antiviral targets for hepatitis D virus infection. Antiviral Res 2017 May 01;141:116-123.

[S3] Pereira IVA, et al. Primary Biliary Acids Inhibit Hepatitis D Virus (HDV) Entry into Human Hepatoma Cells Expressing the Sodium-Taurocholate Cotransporting Polypeptide (NTCP). PLoS One. 2015; 10(2): e0117152.

[S4] Livak KJ, Schmittgen TD. Analysis of relative gene expression data using real-time quantitative PCR and the 2(-Delta Delta C(T)) Method. Methods 2001 Dec;25(4):402-8. doi: 10.1006/meth.2001.1262.

**SUPPLEMENTARY FIGURES**

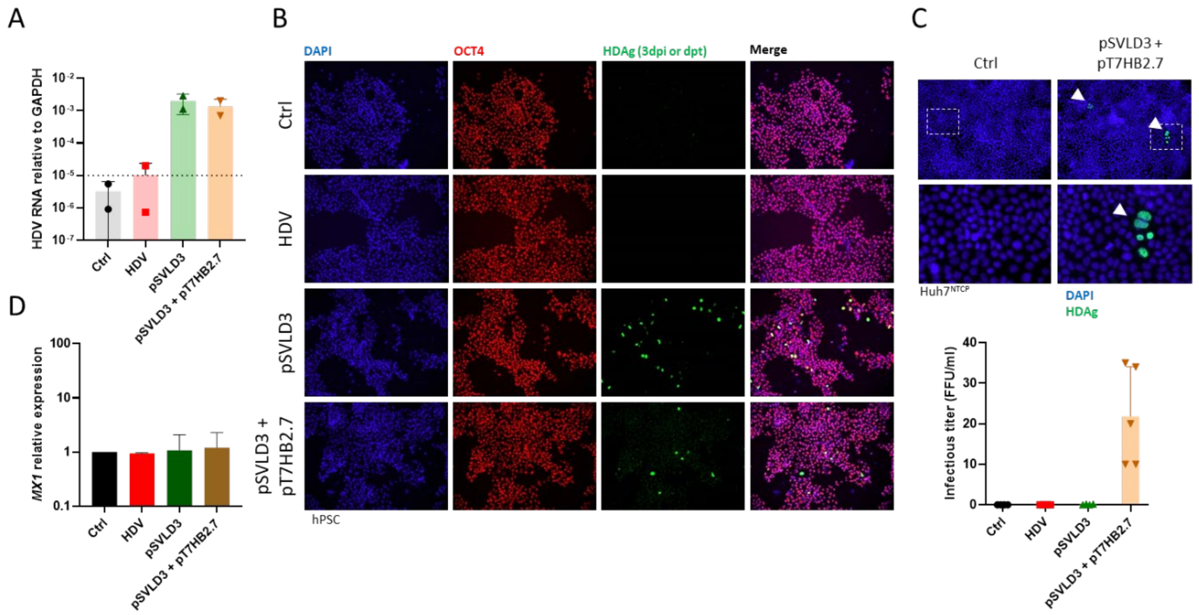


**Supplementary Figure 1: HDV infection or HDV genome transfection of hPSCs.** hPSCs inoculated with HDV, or transfected using Fugene® with the plasmid pSVLD3 (containing a trimer of the genotype 1 HDV genome), without or with the plasmid pT7HB2.7 (encoding the HBV surface antigens). 3 days post inoculation or transfection, (A) intracellular HDV RNA and (B) protein expression of the HDAg were assessed by RTqPCR and IFA (Magnification 10x). (C) Supernatants were collected and transferred on highly permissive Huh7$^{NTCP}$ to assess production of infectious progeny virions, assessed by IFA (Magnification 10x and 40x) and counting of HDAg foci. (D) Induction of the canonical ISGs *MX1* was investigated by RTqPCR.

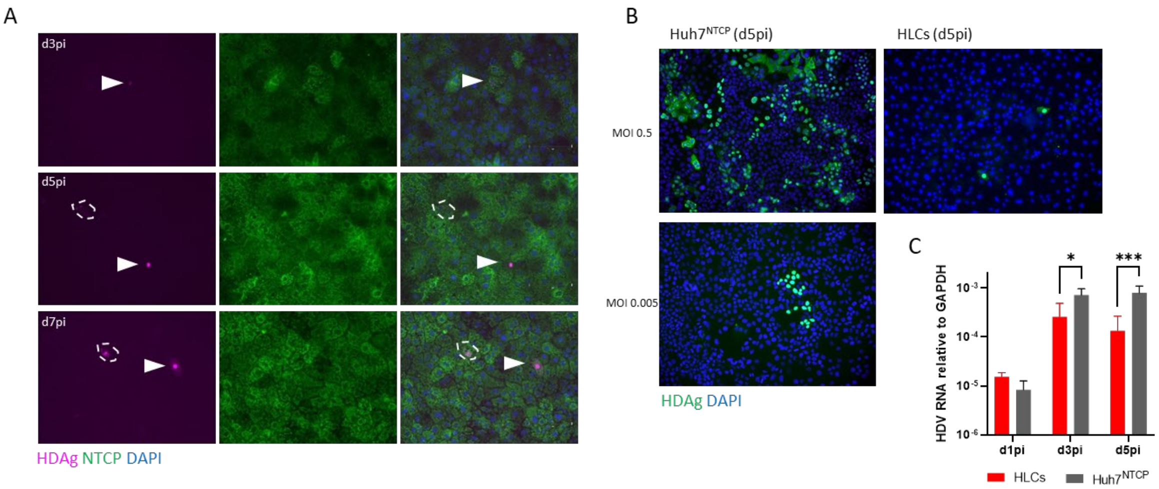


**Supplementary Figure 2: HDAg staining in infected HLCs and comparison to Huh7$^{NTCP}$.** (A) HDAg and NTCP staining of inoculated HLCs (MOI 0.5) at days 3, 5 and 7 pi, assessed by IFA. (B) Typical HDAg staining at d5pi: foci of positive cells in dividing Huh7 vs. single positive cells in quiescent HLCs. (C) Intracellular HDV RNA in HLCs and Huh7$^{NTCP}$ inoculated with HDV at MOI 0.5 and assessed by RTqPCR at d1, 3 and 5pi.

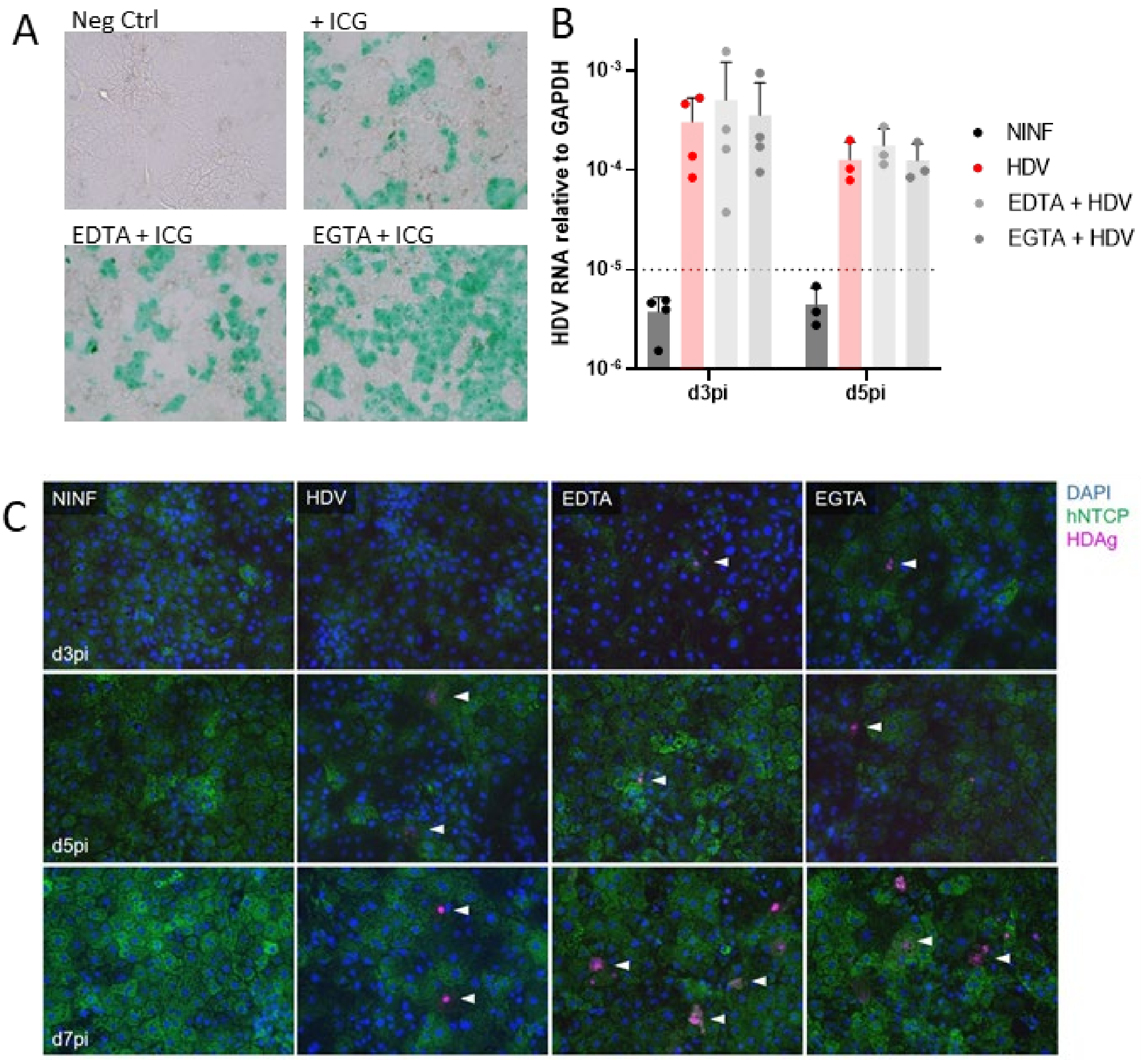


**Supplementary Figure 3: Disruption of tight junctions by EDTA and EGTA does not improve HLCs susceptibility to HDV.** (A) HLCs, control untreated or EDTA/EGTA-treated, were then cultured in presence of 1 mg/mL of Indocyanin Green for 1 hour, and after rinse, NTCP-mediated internalisation was visualised by bright field microscopy. (B) HLCs, control untreated or EDTA/EGTA-treated were then inoculated with HDV as previously described, and intracellular HDV RNA was assessed by RTqPCR at d3 and d5pi. (C) Protein expression and accumulation of HDAg and expression of NTCP were assessed at d3 and d5pi by IFA in these cells. All pictures taken at magnification 10x.

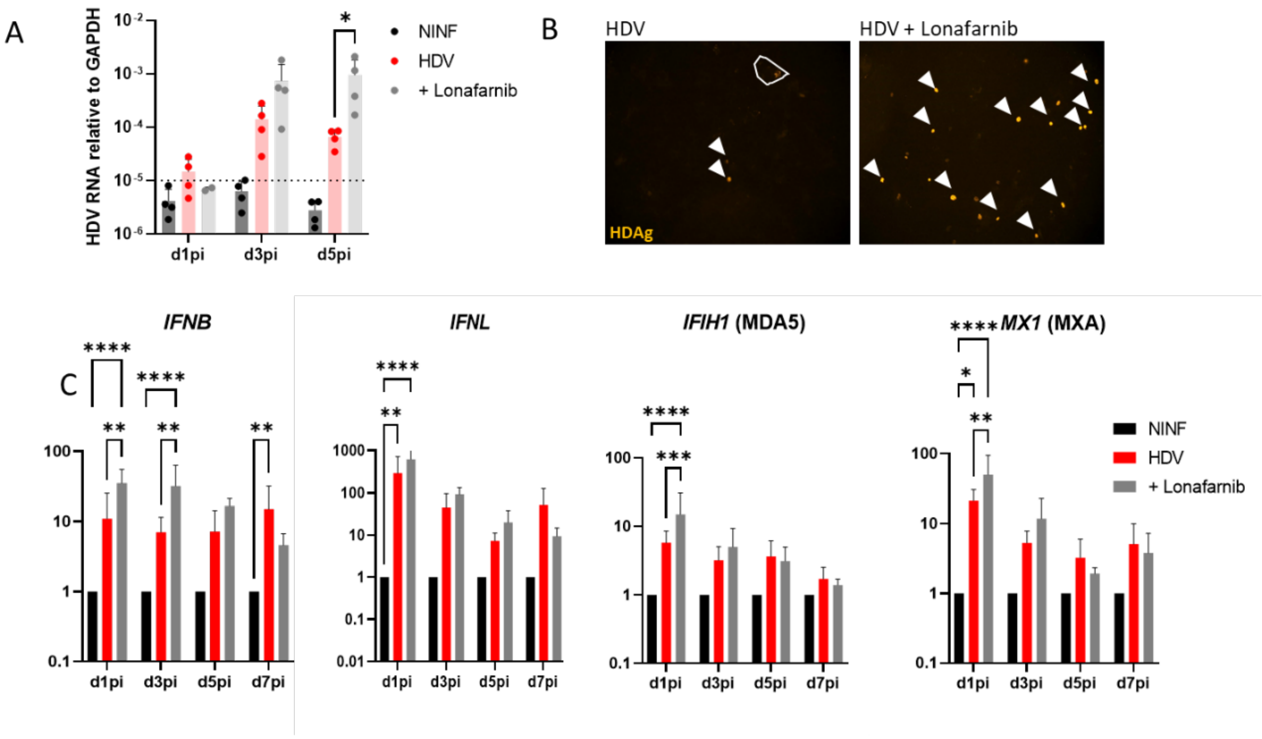


**Supplementary Figure 4: Lonafarnib treatment increases HDV RNA replication and innate immune activation in HLCs.** (A) HLCs, control untreated or treated with 0.1µM of the HDV secretion inhibitor Lonafarnib, were inoculated with HDV as previously described and intracellular HDV RNA was assessed by RTqPCR every 2 days post inoculation. Dots represent intracellular HDV titers of n=4 individual experiments. (B) Protein expression and accumulation of HDAg was assessed 5 dpi by IFA (Magnification 10x) in these cells. (C) Induction of IFNs and ISGs was also assessed upon HDV inoculation in absence or presence of Lonafarnib, expressed as average induction of n=4 individual experiments, relatively to control non infected HLCs (NINF) at the same time points.

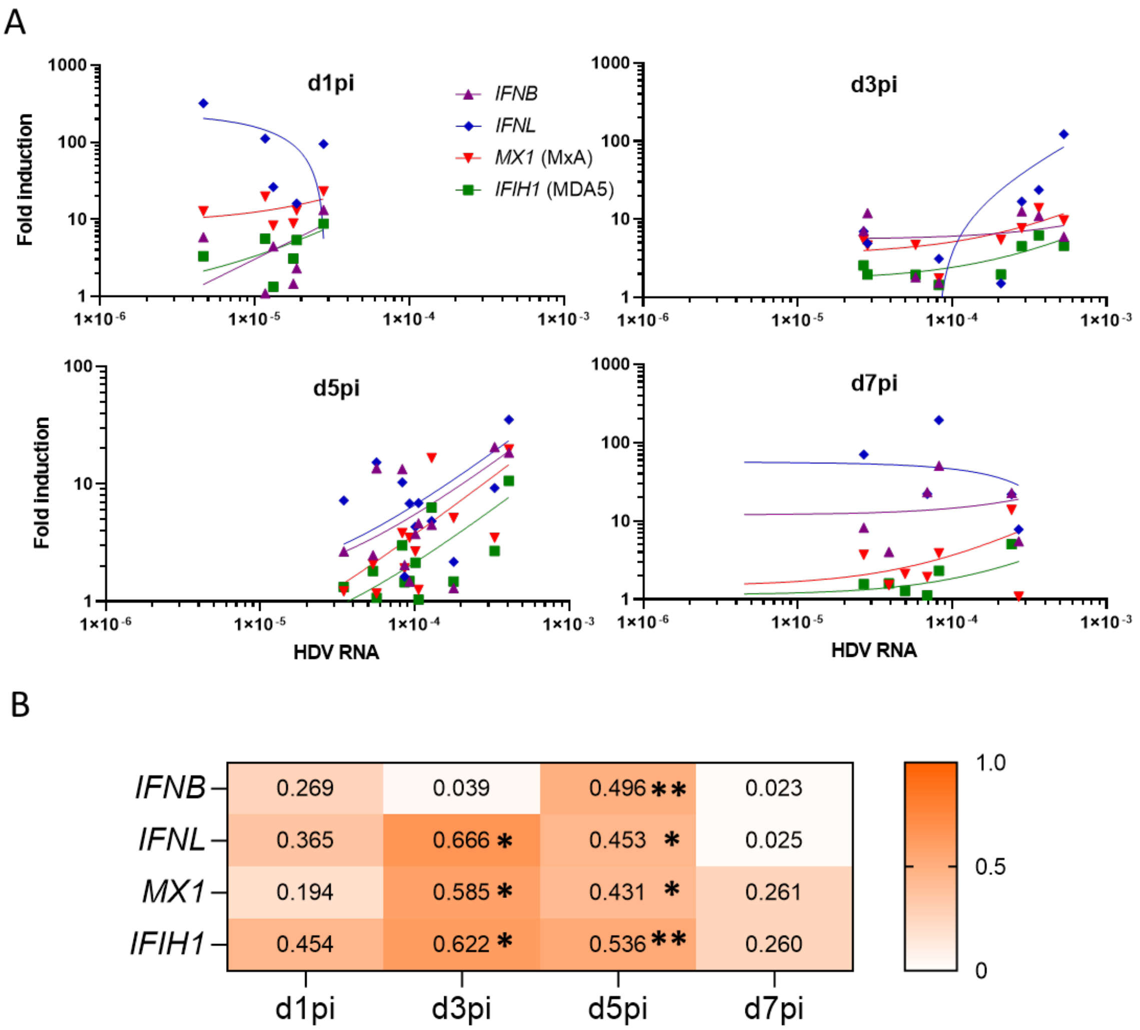


**Supplementary Figure 5: Innate immune activation correlates with level of intracellular HDV RNA.** (A) At the different days post infection, level of intracellular HDV RNA (normalised to GAPDH) was compared to level of induction of *IFNB, IFNL, MX1* and *IFIH1.* Data from 7-12 independent experiments. (B) Heatmap depicts the $R^2$ value of the Pearson's r correlation test for the 16 data sets represented in panel A. Asterisks represent the statistical significance of the Pearson's r correlation test.

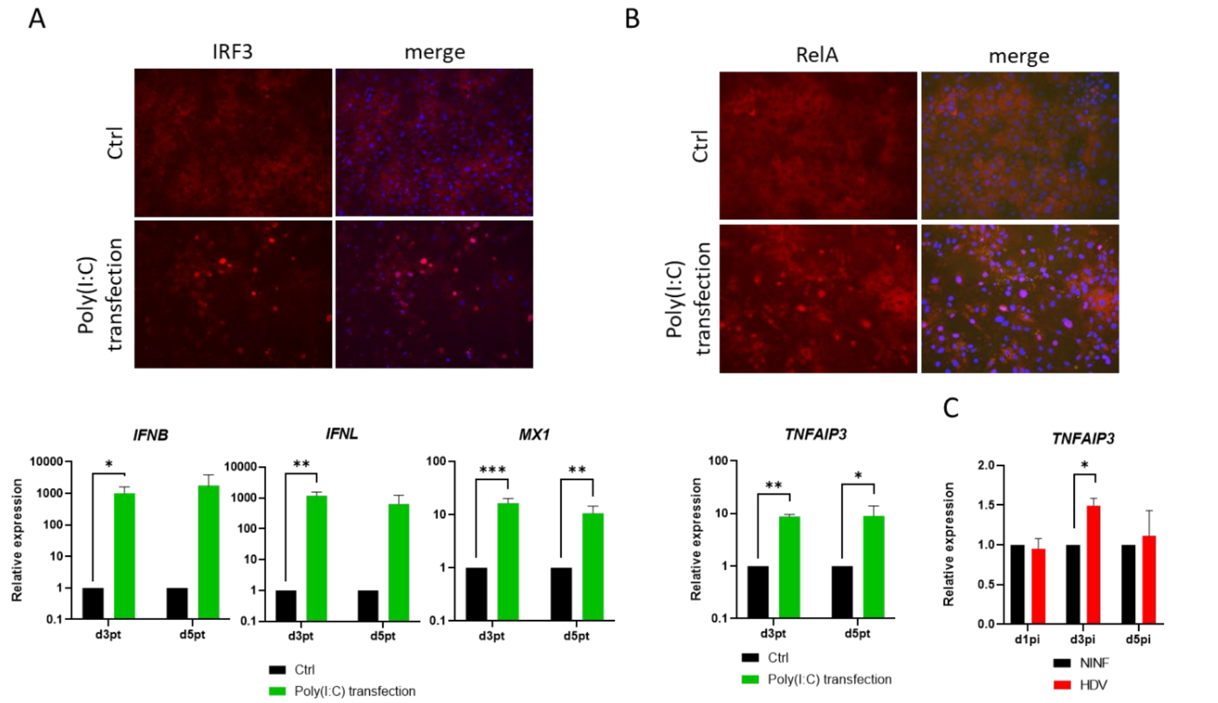


**Supplementary Figure 6: Poly(I:C) transfection activates IFN and NFκB pathways in HLCs.** (A) Nuclear translocation of IRF3 (Magnification 10x) and induction of IFNs and canonical ISG *MX1* (MXA) compared to control untreated cells (Ctrl), assessed by RTqPCR, in HLCs transfected with 1µg/mL Poly(I:C). (B) Nuclear translocation of RelA and induction of NFκB target gene *TNFAIP3* (A20)) upon transfection of HLCs with 1µg/mL Poly(I:C). (C) Induction of NFκB target gene *TNFAIP3* (A20) upon HDV infection of HLCs.

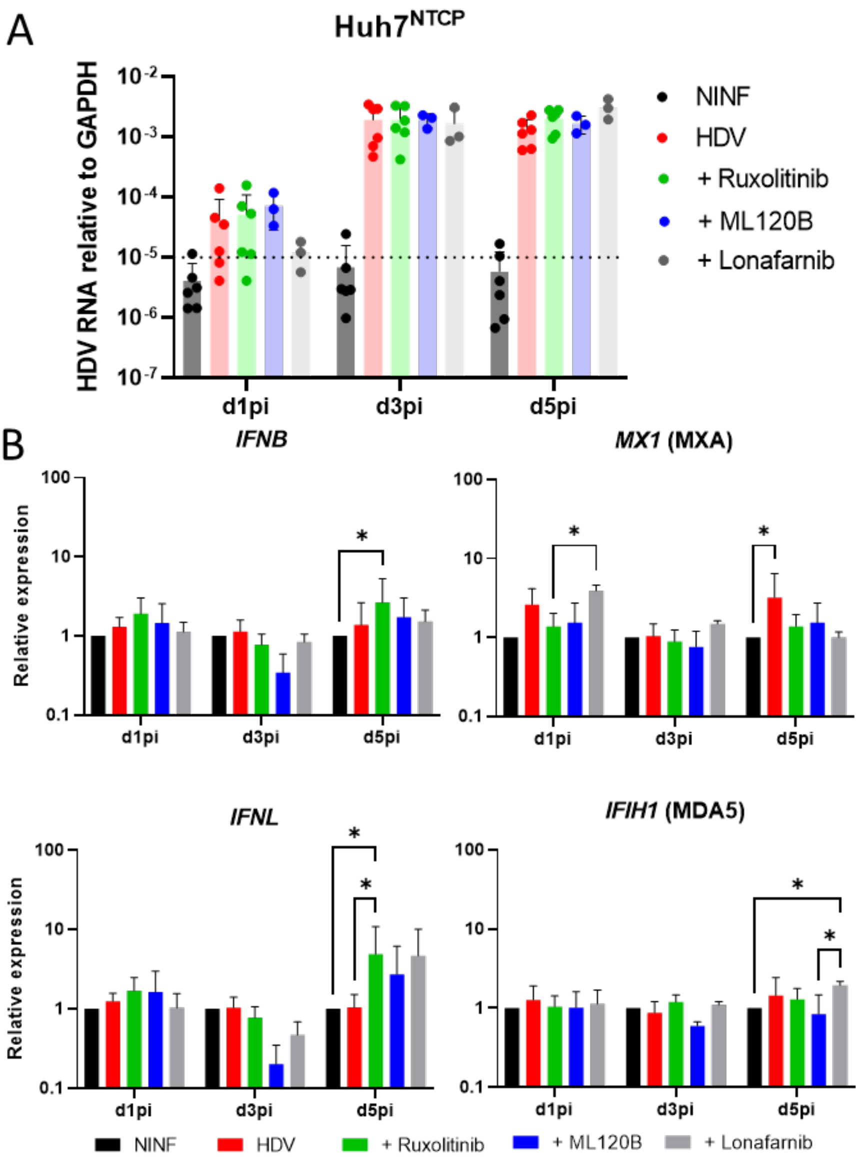


**Supplementary Figure 7: HDV infection does not trigger innate immune activation in** (A-B) Huh7$^{NTCP}$ (n=6) were inoculated with HDV, in absence or presence of the JAK/STAT inhibitor Ruxolitinib, the NFkB inhibitor ML120B, or the assembly inhibitor Lonafarnib. (A) Intracellular HDV RNA and (B) IFNs and ISGs activation were then assessed by RTqPCR at day 1, 3, and 5 post inoculation, expressed relatively to control non infected HLCs (NINF) at similar time points.

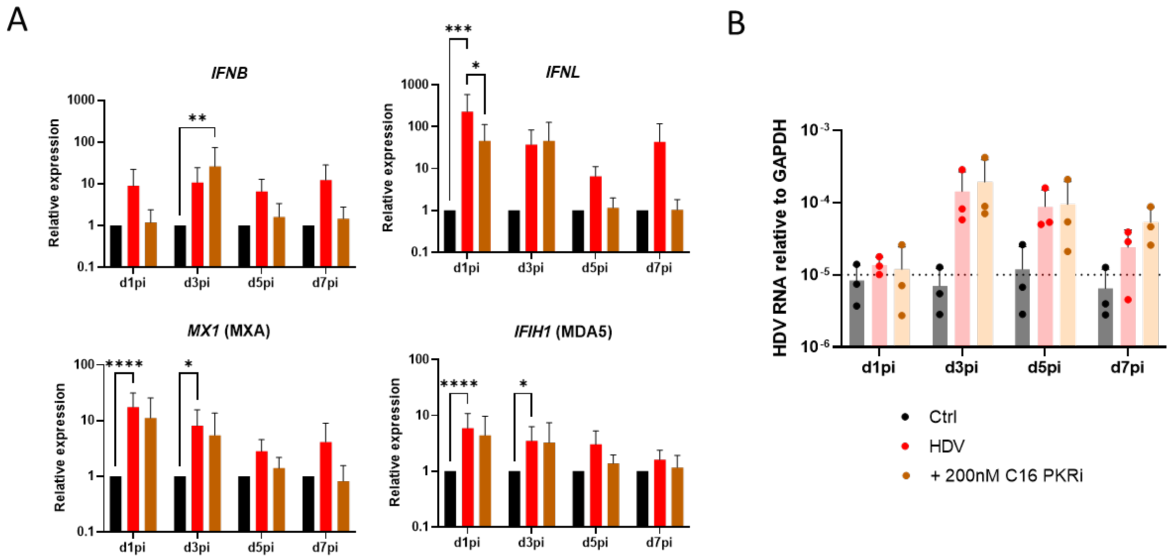


**Supplementary Figure 8: PKR activation does not affect HDV infection of HLCs.** HLCs, control untreated or treated with 200nM of the PKR inhibitor C16, were inoculated with HDV as previously described (n=3). (A) Induction of IFNs and ISGs was assessed upon HDV inoculation in absence or presence of C16, assessed by RTqPCR and expressed relative to control untreated non-infected HLCs (Ctrl). (B) Intracellular HDV RNA was assessed by RTqPCR every 2 days post inoculation, and expressed relative to GAPDH.

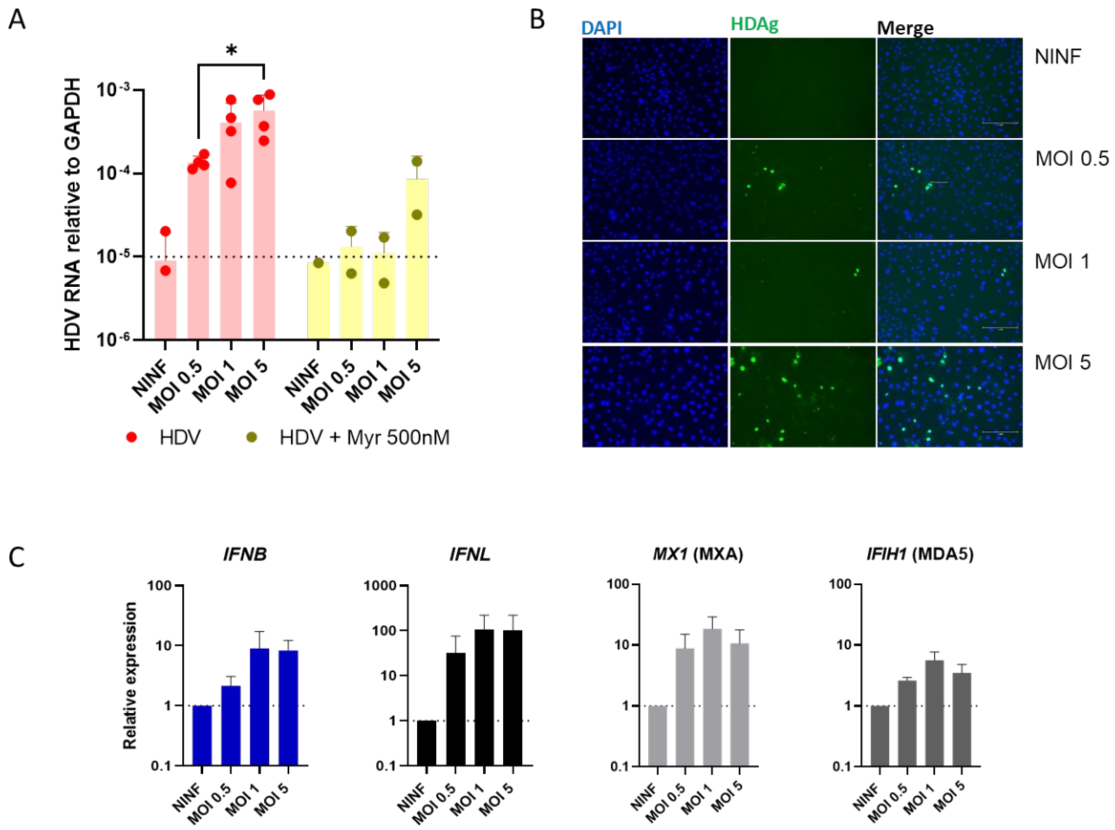


**Supplementary Figure 9: HLCs infected with increasing Multiplicity of Infection (MOI)**. HLCs infection was studied 3 days post inoculation with MOI 0.5, 1 and 5, and compared to control non-inoculated cells (NINF). (A) Intracellular HDV RNA was assessed by RTqPCR, and normalised to GAPDH, in HLCs inoculated in absence or presence of Myrcludex. (B) HDAg expression and percentage of positive cells was assessed by IFA (Magnification 10x). (C) Induction of IFNs and ISGs was assessed by RTqPCR and normalised to control HLCs.
